# Nuclear Speckle Dynamics are Controlled by Polyphosphate Inhibition of CLK Proteins

**DOI:** 10.1101/2025.01.15.633116

**Authors:** Blanca Lázaro, Francisco J. Tadeo-Masa, Andrea Rodriguez, Lucia Ayuso, Joan M Martínez-Láinez, Eva Quandt, Maribel Bernard, Filipy Borghi, Adolfo Saiardi, Jonàs Juan-Mateu, Javier Jiménez, Josep Clotet, Samuel Bru

## Abstract

Nuclear speckles (NS) are membrane-less nuclear organelles that act as critical hubs for pre-mRNA splicing. Defects in splicing are linked to several human diseases, including cancer, Alzheimer’s disease, and dystrophies. While CLK kinases regulate the mobilization of splicing factors from NS, the molecular mechanisms underlying NS assembly and dissolution remain unclear. Using an adaptation of the Biotinylation by Antibody Recognition (BAR) technique, we identify polyphosphate (polyP) as a novel and essential regulator of NS dynamics. Polyphosphate, a highly conserved polyanion composed of a chain of phosphate molecules, is involved in several functions in mammalian cells. Here, we show that polyP interacts with the NS core component SRRM2, and polyP depletion disrupts NS organization releasing splicing factors into the nucleoplasm. RNA-seq analysis reveals that polyP depletion increases exon inclusion, particularly in long genes with multiple exons, highlighting its role in splicing regulation. Mechanistically, we demonstrate that polyP acts as a physiological inhibitor of CLK3 kinase, preventing the phosphorylation of SR proteins and thereby maintaining NS stability. Our findings not only expand our understanding of NS biology but also provide new insights into the polyP involvement in splicing-related diseases.

**GRAPHICAL ABSTRACT:** 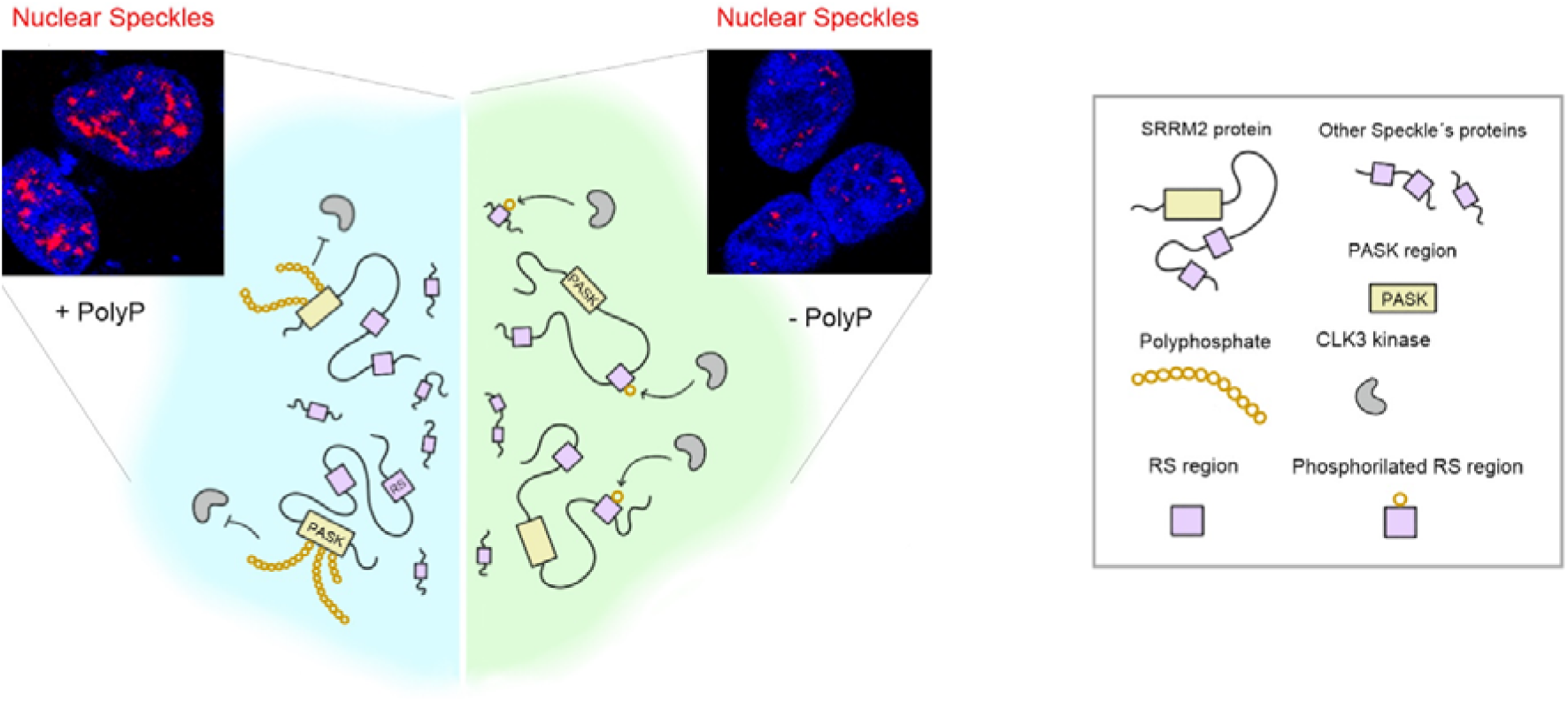

## INTRODUCTION

Initially shown by Ramón y Cajal (1), nucleoplasm is compartmentalized in several types of membrane-less nuclear bodies whose main role is concentrating the biochemical machinery required for efficiently performing a function (2). Specific proteins and polyanionic molecules such as RNAs, and DNA can gather in the nucleoplasm to form the nucleolus, Cajal Bodies, PML bodies, speckles, and others (3). Particularly, nuclear speckles (NS) are membrane-less bodies enriched in pre-mRNA splicing factors in the interchromatin areas of the mammalian cell’s nucleoplasm (4). NS play a crucial role in buffering the nucleoplasmatic concentration of different splicing factors, thereby influencing splicing efficiency and the alternative splicing of nascent RNAs throughout the nucleus (5). NS are also associated with active transcription sites to enhance the transcription of proximal genes (6–8). Biochemical purification methods (9) and, more recently, high-throughput screens (10, 11) have produced several catalogs of NS resident proteins. Among these, SRRM2 and SON stand out as proteins of particular interest due to their essential role in maintaining NS assembly (12). However, the depletion of either SRRM2 or SON alone cannot fully disrupt NS formation. This suggests that both proteins function as central nodes within the NS network, acting together to preserve their integrity (13).

Several sequence elements have been proposed to push proteins into NS, including poly-histidine stretches (13, 14) and arginine-serine (RS) repeats (15). More recently, other sequence elements have been proposed such as mixed-charged domains (MCDs), particularly stretches of arginine (but not lysine) and glutamic acid residues within low-complexity regions (16). Interestingly, both SRRM2 and SON contain RS domains embedded within low-complexity regions, which are essential for the formation of higher-order oligomers triggering NS condensation (13).

The RS domains undergo dynamic cycles of phosphorylation and dephosphorylation, and the degree of negative charge on these domains can influence the aggregation of splicing factors (17). Specifically, increasing the negative charge (phosphorylation) facilitates the splicing factor’s mobility and the disassembly of the NS (16). Accordingly, the expression of dual-specificity kinases, such as CLK1 (18) and DYRK3 (19), triggers a phosphorylation-dependent disassembly of splicing speckles; while the overexpression of PP1 phosphatases increases NS cohesion and mRNA enrichment (20). These findings underscore the pivotal role of kinases and phosphatases in regulating the physiology of NS and splicing processes.

Alternative splicing is an essential co-transcriptional process that allows the exons of a gene to join in different combinations resulting in distinct but related mRNA transcripts. The splicing of pre-mRNAs derived from small genes, which contain short introns and a low percentage of GC content, occurs near NS (21). In contrast, pre-mRNAs from large genes, with long introns and a low percentage of GC content, are spliced farther away from the NS, requiring the mobilization of splicing factors from NS (21, 22). Additionally, the presence of free splicing factors in the nucleoplasm has been associated with a faster release of the mRNA from the gene (18). This indicates that cells can regulate gene expression by modulating the availability of splicing factors.

Polyphosphate (polyP) is a polymer of a varying number of ortho-phosphates (from a few to several hundred) linked by high-energy bonds (23) and, in consequence, a highly negatively charged molecule, a polyanion like RNA and DNA. Polyphosphate is an intriguing cellular element present in all cells from archaea to humans and involved in a plethora of physiological functions. Indeed, human polyP functions in blood coagulation (24), inflammation (25), stress response (26), mitochondria and energetic metabolism (27), cell cycle (28), bone formation (29) and protein folding (30). In addition, polyP levels are altered in pathological conditions such as amyloidogenic diseases (23, 31), other neurodegenerative diseases such as ALS and FTD (32), and proliferative maladies (33–35). A proposed mechanism for regulating the diverse array of polyP cellular functions involves the poly-phosphorylation of lysine residues embedded in regions enriched with aspartic acid, glutamic acid, and serine—referred to as PASK (Poly-Acidic-Serine-Lysine) domains (36). Another suggested mechanism is polyP specific interaction with lysine stretches (37). Additionally, polyP has been shown to inhibit DYRK1 kinase activity through selective binding to histidine repeats (38). Further evidence will help in establishing whether the polyP interaction with PASK, Lys and His regions represents the general mechanism by which polyP modulates cellular processes in mammals.

Metazoan polyP is a highly dynamic molecule with a high turnover rate (39) prominently localized in the nucleus(39–41). Recently, immunolocalization methods using the PPBD peptide (<u>P</u>px1 <u>P</u>olyP <u>B</u>inding <u>D</u>omain) which specifically binds to polyP (42, 43), show a characteristic punctuated nuclear signal suggesting that polyP might localize within some of the membrane-less nuclear bodies (43, 44).

Identifying interactors under the rationale of ‘guilty by association’ (45) remains an interesting strategy for characterizing the biological functions of poorly understood molecules. Over the last decade, Proximity Labeling has enabled the dissection of complex interactomes with remarkable spatial and temporal resolution (46). To overcome some limitations of traditional Proximity Labeling methods, the BAR (Biotinylation by Antibody Recognition) method was developed, which replaces fusion enzymes with horseradish peroxidase (HRP)-conjugated antibodies (47). The BAR method directs biotinylation to neighboring proteins in fixed cells, enabling the detection of transient interactions of high-turnover molecules within their native spatial context, interactions that are often challenging to capture with conventional protein interaction techniques.

Here, we provide a solid workflow to target polyP proximal proteome (PPP) using an adaptation of the BAR technique. We have taken advantage of the PPBD peptide to target polyP in fixed cells and drive the biotinylation of its proximal proteins. We found that polyP is an unexpected element of the NS that maintains its integrity and functionality by restricting the action of CLK kinases.

## MATERIAL AND METHODS

### Cell lines and culture conditions

All experiments were performed using HEK293T cell line. Cells were grown in Dulbeccós Modified Eagle Medium (Sigma, D5671) supplemented with 10% fetal bovine serum (Labclinics, FBS-12A), 1% glutamax (Labclinics, X0551-100)) and 1% penicillin/streptomycin (Sigma, P0781-100). Cells were grown in humidified air at 37°C and 5% CO_2_ atmosphere. Mycoplasma contamination was monitored periodically. In some experiments, the cells were treated with doxycycline (Sigma, D1822), Menadione (Sigma, M5625) or Hexanediol (Sigma, H11807).

### Transfection

FuGENE (Promega, E2311) system was used to transfect DNA plasmids. The protocol was as indicated by the purchaser. Briefly, 25 µL of the transfecting mix was added to every p24 plate well. Transfecting mix includes 250 ng of the plasmid and 1 µL of FuGENE reagent in 25 µL of culture medium without FBS and penicillin-streptomycin. The mix was incubated at RT for 5 min and then was added to the cells. For larger wells, the volumes were scaled up. Lipofectamine 2000 (ThermoFisher, 11668019) system was used to transfect siRNAs. The protocol was as indicated by the purchaser. Briefly, 100 µL of the transfecting mix was added to every p24 plate well. Transfecting mix includes 5 pmol siRNA oligomer (10 nM final concentration) in 50 µL of culture medium without FBS and penicillin-streptomycin and 50 µL of lipofectamine mix (0.8 µL of lipofectamine 2000 in 50 µL of culture medium without FBS and antibiotics). Both components, previously kept at RT for 5 min, were mixed gently and incubated for 20 min at RT. Once the lipofectamine-siRNA was added to the cells, the plates were gently agitated and incubated at 37 °C in a 5% CO_2_ containing atmosphere for 24-48 h. For larger wells, the volumes were scaled up. The sequence of the siRNAs used is:

Scrambled (IDT, 51-01-14-04); NUDT3 (rGrCrArCrArGrGrArCrGrUrArUrGrUrCrUrArUrGrUGC); SRRM2 (ThermoFisher, s24004); SON (ThermoFisher, s13278) and SRRM1 (ThermoFisher, s20020).

### Plasmids and recombinant proteins

The plasmids are in Table 1. *E. coli BL21-CodonPlus(DE3)-RIPL* was used for expression of recombinant proteins. Proteins were induced by adding 1 mM IPTG to a culture in exponential phase and incubated for 4 h at 30°C. Cells were lysed in lysis buffer (25 mM NaH2PO4, 10 mM imidazole, 1 % triton, 2 mM β-mercaptoethanol, 150 mM NaCl, protease inhibitors (ThermoFisher, A322955), 0.4 mg/mL lysozyme, 4 U/mL benzonase). Proteins were purified using a AKTA (Cytiva, 29022094). Proteins were quantified by Bradford and purity was check by PAGE and Coomassie staining (BioRad, 1610787).

**Table 1:** Plasmids.

| Plasmid | Commercial supplier | Reference |
| --- | --- | --- |
| pWPI-NUDT3-flag | Our lab. | N/A |
| pWPI-NLS-Ppx1-eGFP-FLAG | Our lab. | N/A |
| pRetroX-Tet-on-Advanced | Takara | 632104 |
| pRetroX-Tight-Pur | Takara | 632105 |
| pRetroX-Tight-Pur-NUDT3 | Our lab. | N/A |
| pX459 | Addgene | 62988 |
| pTRCHisB-PASK<sub>SRRM2</sub>-mCherry | Our lab. | N/A |
| pTRCHisB-RS<sub>SRRM2</sub>-mCherry | Our lab. | N/A |
| pTRCHisB-mCherry | Our lab. | N/A |
| CMV-LUC2CP-intron-ARE | Addgene | 62858 |
| CMV-LUC2CP-ARE | Addgene | 62857 |

### Western and dot blots

For western blot, cells were scrapped in lysis buffer (20 mM Tris-HCl pH 7.4, 5 mM EDTA, 1% NP40 and 150 mM NaCl) containing protease and phosphatases (when necessary) inhibitor cocktails (ThermoFisher, A32955 and ThermoFisher, A32957 respectively). The lysate was centrifugated and the protein concentration was quantified using Bradford (BioRad, 5000006). Samples were denatured at 95 °C for 5 min and 30 µg of total protein was separated in 4-15% SDS-PAGE gels (BioRad, 5671084) and transferred to PVDF membranes (Sigma, IPVH00010). Membranes were incubated in blocking buffer (TBS-T containing 5% non-fat milk) for 30 min at RT and incubated with the primary antibody (Table 2) in blocking buffer O/N at 4 °C. After washing, membranes were incubated with horseradish peroxidase (HRP) conjugated anti-mouse or -rabbit IgG secondary antibodies for 1 h at RT. After additional washes, the membranes were developed using Luminata Forte western HRP substrate (Sigma, WBLUF0500) following manufacturer’s instructions. Images were taken using the ChemiDoc Imaging System (BioRad) and analyzed with Fiji software (48).

**Table 2:** antibodies.

| Antibody | Commercial supplier | Dilution |
| --- | --- | --- |
| Anti-NUDT3 | Sigma (SAB2700523) | WB 1:2000 |
| Anti-FLAG | Sigma (F3165) | WB 1:1000 |
| Anti-Biotin | Sigma (B3640) | WB 1:1000 |
| Streptavidin-AF488 | ThermoFisher (S11223) | IF 2 µg/mL |
| Anti-SC35-AF647* | Santa Cruz (sc-53518 AF647) | IF 1:50 |
| Anti-SON-AF647 | Santa Cruz (sc-398508) | IF 1:200 |
| Anti-SRRM1 | Abcam (ab221061) | IF 1:500 |
| Anti-RBM25 | Sigma (HPA070713-100UL) | IF 1:250 |
| Anti-Xpress | ThermoFisher (R91025) | IF 5 µg/mL |
| Anti-PML-AF647 | Santa Cruz (sc-966) | IF 1:500 |
| Anti-Coilin-AF647 | Santa Cruz (sc-56298) | IF 1:100 |
| Anti-Fibrillarin | Abcam (ab5821) | IF 1:100 |
| Anti-H3K9me3 | Abcam (ab8898) | IF 1:50 |
| Alexa Fluor 488 conjugated goat anti mouse IgG | ThermoFisher (A11001) | IF 1:1000 |
| Alexa Fluor 594 conjugated goat anti mouse IgG | ThermoFisher (A11005) | IF 1:1000 |
| Alexa Fluor 633 conjugated goat anti mouse IgG | ThermoFisher (A21052) | IF 1:1000 |
| Alexa Fluor 488 goat anti rabbit IgG | ThermoFisher (A11008) | IF 1:1000 |
| Alexa Fluor 568 goat anti rabbit IgG | ThermoFisher (A11011) | IF 1:1000 |
| Alexa Fluor 647 goat anti rabbit IgG | ThermoFisher (A21244) | IF 1:1000 |
| anti-Penta-His | Qiagen, 34660 | WB 1:1000 |
| anti-pSR | Sigma, MABE50 | WB 1:5000 |
| anti-Thiophosphate ester | Abcam (ab92570) | WB 1:5000 |

In dot-blot assays, PVDF membranes were used to check PIP2 and Nylon Biodyne B membranes (ThermoFisher, 77016) for RNA and polyP. Samples (1-5 µL) were attached to the membrane and allowed to dry. Membranes were blocked with 0.01% TBS; 0.01% Tween-20 and 5% BSA for 1 h at RT. Membranes were incubated in the previous blocking buffer containing the corresponding probe. Fluorescent probes (mCherry, PASK-mCherry or RS-mCherry) were at 2 µM. PPBD was at 10 µg/mL accompanied by α-His. α-PIP2 was at 1:500. Fluorescence imaging for fluorescent probes or secondary antibody incubation were as in western blot.

### Immunofluorescence and confocal microscopy

For immunofluorescence, cells were grown in plate wells containing a poly-lysine treated coverslip (Sigma, P4707). Cells were fixed using PBS containing 10 % paraformaldehyde. Cells were blocked and permeabilized with blocking solution (PBS; 0.5% Tween 20 and 2.5% goat serum) for 1 h at RT. Cells were incubated O/N at 4°C with the specific antibodies in blocking solution (Table 2). For polyP detection, cells were incubated O/N at 4°C with the Xpress-tagged PPBD probe and the anti-Xpress antibody as described in (44). After washing 3 times with PBS, secondary antibodies with different Alexa fluorophores (Invitrogen) were added (1:1000 in blocking solution) for 1 h at RT. Samples were washed 3 times with PBS and Hoechst 33342 (Sigma, 14530) for 5 min at RT (1 µg/mL) was used for nuclei staining. Samples were mounted using Fluoromount-G (Southern Biotech, 0100-01). The microscope was a Leica confocal SP8. Fiji software was used for quantification. ROIs were defined applying a threshold in the Hoechst channel with a previous Gaussian B filter plugin. To ensure accuracy in the ROIs and to create the mask, the plugins: “fill holes” and “watershed” were applied. The ROI mask was overlaid on the channel of interest and the raw integrated density was obtained. The raw integrated density mean in each field was calculated. The Pearson correlation coefficient was calculated using the appropriate plugin in ImageJ program.

### Biotinylation by Antibody Recognition (BAR)

As described by Bar et al. (2018). The kit was Biotin XX Tyramide SuperBoost™ biotin labelling kit (ThermoFisher, B40911) and following manufacturer’s instructions.

Briefly, 2x10^6^ cells were fixed in PBS with 10% PFA, treated with 3% H_2_O_2_ for 10 min at RT (to abrogate endogenous peroxidase activity) and blocked with PBS containing 1% BSA and 0.5% Tween-20 for 1 h at RT. The cells were incubated in blocking buffer O/N at 4°C with anti SRRM2 antibody (SC35 see antibodies table) Alexa 647 conjugated to determine the speckle proteome, or with a complex of PPBD peptide (2 µg/mL) and α-Xpress (4 µg/mL) to determine the polyP interactome. Cells were treated with the kit’s secondary antibody. After washing 3 times with PBS, the biotinylation reaction was carried out using the reaction buffer containing H_2_O_2_ and biotin-tyramide. The reaction was stopped after 3 min (Figure S1-A) with the stop buffer included in the kit. To analyze the biotinylated proteins, cells were scraped into 100 µL of PBS-T (PBS containing 0.1% Tween-20) and lysed by adding 30 µL of 10 % SDS and 20 µL of 10 % sodium deoxycholate and incubating for 1 h at 99°C with agitation. The lysate was centrifuged at maximum speed. The supernatant was brought to 1 mL with PBS-T and incubated with 50 µL of magnetic beads (ThermoFisher, 88816) for 2 h at RT under gentle agitation. Samples were washed once with PBS-T, followed by PBS-T containing 1M NaCl, and finally with PBS. The dried beads were analyzed by Mass Spec to detect the proteins. All the steps were monitored by western blot.

### Protein identification by LC-MS/MS

The proteomic analysis was performed in the Proteomics Unit of Complutense University of Madrid. Briefly, the dried beads containing the biotinylated proteins were digesting using IST kit (Preomics) and following the manufacturer instructions. Peptides (1µg) were analysed by liquid nano-chromatography (Easy-nLC 1000, Thermo Scientific, Bremen, Germany) coupled to a high-resolution mass spectrometer Q-Exactive HF (Thermo Scientific). Peptides were concentrated by reverse phase chromatography using an Acclaim PepMap 100 precolum (Thermo Scientific) and separated in a reverse phase C18 Picofrit analytical column (Thermo Scientific) Peptides were detected in the range of 350-2,000 Da. For the protein identification the peptides data were analysed using Proteome Discoverer 2.5 (Thermo Scientific) software using MASCOT v.28 as the search engine. The database was Uniprot restricted to *homo sapiens* and a contaminants database.

For a hit to be considered a polyP interactor two thresholds were applied. First, the Mascot Score greater than 85. Second, the Mascot score must be (at least) 9 times higher in the experimental sample than in the control. The databases for analyzing the list of interactor were: Localization, the g:Profiler database (version e111_eg58_p18_f463989d), To determine whether the proteins were in the nucleus (GO:0005654) or the cytosol (GO:0005829), as well as their specific location within the nucleus: Nuclear speckle (GO:0016607), Nucleolus (GO:0005730), Nuclear matrix (GO:0016363), Chromatin (GO:0000785), Cajal body (GO:0015030), Nuclear envelope (GO:0005829), Nuclear membrane (GO:0031965), PML body (GO:0016605), and Paraspeckle (GO:0042382). To determine the function, the ShinyGO 0.80 web was used, searching the KEGG database. The list was compared with the following lists in the database: spliceosome (hsa03040), DNA replication (hsa03030), ribosome biogenesis (hsa03008), mRNA surveillance (hsa03015), viral live cycle HIV-1 (hsa03250), nucleocytoplasmic transport (hsa03013), nucleotide excision repair (hsa03420), cell cycle (hsa04110), mismatch repair (hsa03430), RNA polymerase (hsa03020), homologous recombination (hsa03440), base excision repair (hsa03410), Fanconi anemia (hsa03460) and cytosolic DNA-sensing (hsa04623). To determine the protein domains, the MEME web tool (https://meme-suite.org/meme/tools/meme) version 5.5.5 was used. We compared the polyP-interacting proteins with the domains in the Interpro database, obtaining a False Discovery Rate (FDR) from each comparison, which showed significance for several domain families.

### Differential splicing events determination by RNA seq analysis

Cells were transfected with either pWPI empty, pWPI-Ppx1, siRNA scramble, or siSRRM2. After 48 h, total RNA was isolated using the TRIzol (ThermoFisher) extraction protocol provided by supplier. RNA sequencing was performed by Macrogen (https://dna.macrogen.com/) using a NovaSeq6000 platform and generating 100 million reads (150 nt paired-end) per sample.

### Electrophoretic mobility shift assay (EMSA)

10 µg of the corresponding peptide was mixed with different amounts of commercial polyP water. The samples were incubated for 1 h at RT and then separated in a 1% agarose gel in TBE buffer for 1 h at 100V. The mobility was assessed by the red fluorescence. The RBD domain (49) is defined between residues 186 and 246 of SRRM2. The UPR used was three repeats of the GRSRSRTPA sequence.

### Splicing luciferase assay

The reporter system was as described in (50) with some modifications. Briefly, cells were grown in 6-well dishes and transfected 24 h later with empty pWPI, pWPI-NUDT3, or pWPI-Ppx1. Cells were reseeded in a p96 plate and after 24 h cells were transfected with the luciferase reporter plasmid CMV-LUC2CP/intron or control CMV-LUC2CP (Table 1). After 72 h, D-luciferin (Sigma Aldrich 2591-17-5) was added to a final concentration of 1 mg/mL. Luciferase kinetics were measured at 37 °C using a Synergy HT plate reader the maximum luminescence value was considered (aprox. 1-2 min after adding luciferin). Isoginkgetin (Sigma Aldrich, 416154) was added at 35 µM.

### *In vitro* phosphorylation assay

GST-CLK3 (0.12 µM), either purified from *E. coli* or from insect cells (SinoBiological, C60-10G-10) was added with either GST-SRx4 or mCherry-UPRx3 (1.6 µM) to kinase buffer (50 mM HEPES pH 7.5, 150 mM NaCl, 10 mM MgCl_2_, 5 mM EGTA, 1mM DTT and 1 mM ATP) to a final volume of 20 µL. Polyphosphate P14 or P130 was added. The reaction mix was incubated at 30 ⁰C for 45 min. Samples were subjected to SDS/ PAGE and probed for SR-phosphorylation.

In the CDK16 kinase assay, GST-CDK16 (0.2 μM) and GST-CCNY (0.4 μM) were mixed in a kinase buffer (500 mM HEPES pH 7.5, 500 mM NaCl, 1 mM MgCl_2_, 50 mM EGTA, 10 mM DTT) and 1 mM ATP-γS (Axxora, Farmingdale, NY, USA) were mixed ant incubated at 30 °C for 45 min. Reactions were quenched by adding 50 mM EDTA and incubated with 0.5 mg/mL para-nitro-benzo-mesylate (PNBM), kindly provided by K. M. Shokat, at 25 °C for 45 min. PNBM samples were subjected to SDS/ PAGE and probed for thio-phosphorylation with specific antibodies.

## RESULTS

### Proximity biotinylation of polyP nuclear depots reveals its localization in the nuclear speckle

A strategy to unveil the role of polyP in human cells is to assess its partners. We adapted the Biotinylation by Antibody Recognition (BAR) method (47) using PPBD peptide to bind polyP and label the proteins in its vicinity as an approach for obtaining the polyP proximal proteome (PPP) (Fig. 1A). Considering that formaldehyde fixation does not affect polyP structure but rapidly stops the biochemical machinery of the cell, it will maximize the polyP detection and enable the identification of its proximity proteome by mass spectrometry. After setting up for reducing noise and false positives (Fig S1A), the biotinylated proteins nicely colocalize with the immunolocalized polyP (Fig. 1B). Interestingly, both the polyP and biotinylated proteins are found in areas poorly stained with Hoechst (Fig. 1C) proposed as regions of intense transcription. Biotin detection shows that only biotinylated proteins are obtained when the polyP-binding PPBD is present, indicating the high specificity of the assay (Fig. S1B). Mass spec analysis reveals that 10 out of the top 22 PPP proteins (highlighted in red) with the best comparative Mascot Score have been previously identified as NS residents (Fig. 1D). Notably, and the scaffold proteins SON and SRRM2, which are essential for maintaining NS integrity (12), are among the top PPPs hits. The complete list of the 285 PPP proteins is in supplemental Table 1. GO analysis of the PPP shows a clear nucleoplasm localization (Fig. 1E) and enrichment in the NS (Fig. 1F). Looking at the protein functions, splicing first, and then RNA metabolism proteins, are the most prominent (Fig. 1G). This is confirmed by a domain-containing analysis showing the presence of RNA recognition domains, helicase domains, and well-recognized polyanion interacting domains (Fig. 1H). Confirming the analysis presented above, the polyP immunolocalization mostly colocalizes with SRRM2 a well-recognized NS protein (Fig. 1I), with a Pearson correlation coefficient of 0.8 (Fig. 1J) much higher than the correlation coefficient for other nuclear organelles markers. SRRM2 and polyP are again excluded from the nucleus areas with brighter Hoechst staining (Fig. S1C). Finally, a second BAR analysis using an SRRM2-specific antibody, which aligns with previous NS proteomic studies ((9–11), Fig. S1D and S1E), reveals that 76% of SRRM2-interacting proteins are included in the PPP (Fig. 1K). This confirms that the PPP largely comprises NS proteins, reinforcing the notion that polyP is a new component of the NS.

**Figure 1.**
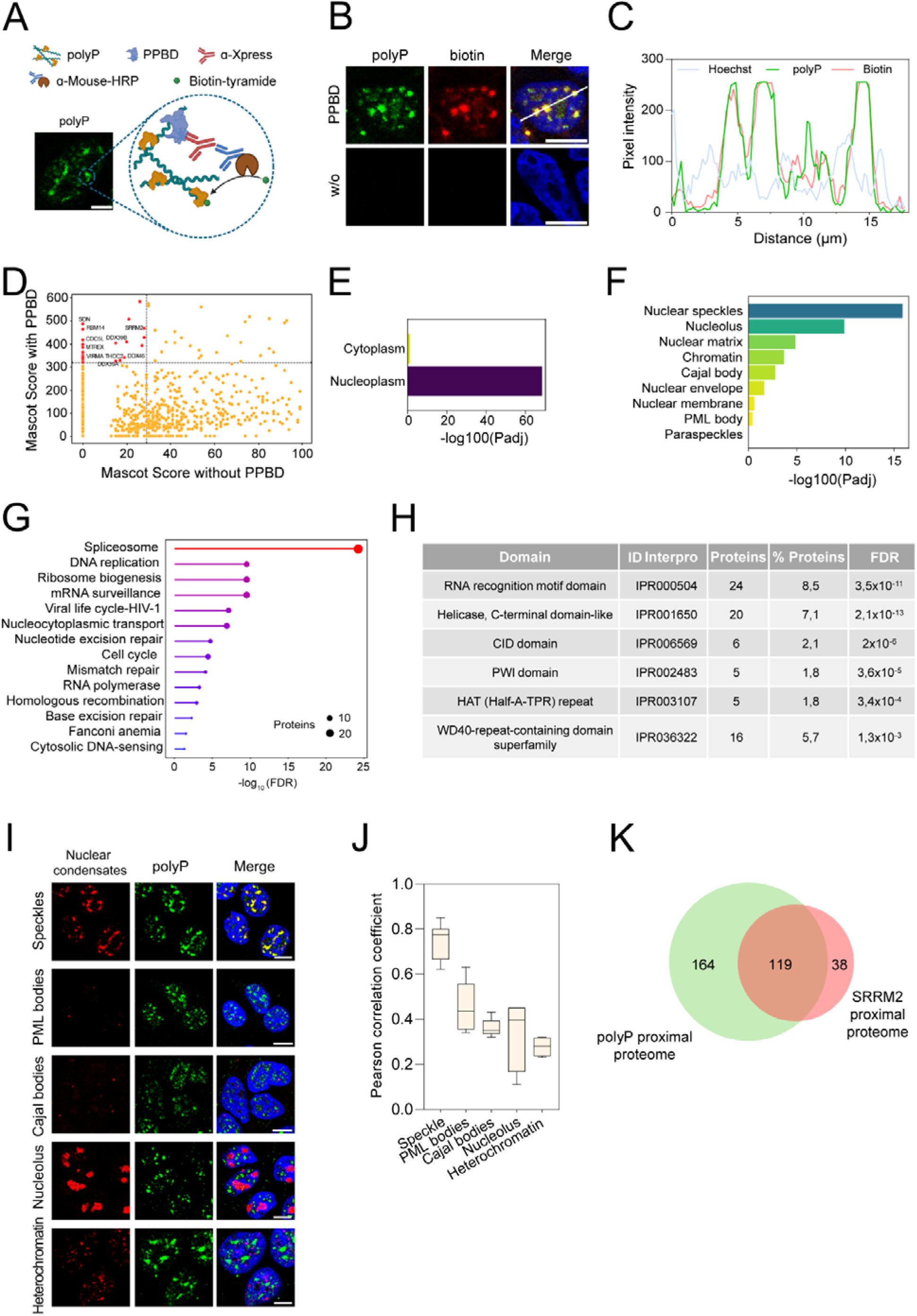
Polyphosphate is primarily found in NS and associated with splicing proteins. (A) PolyP interactome labeling strategy using proximity labeling technique (BAR). The scale bar represents 5 µm. (B) Confocal images of polyP (using PPBD) and associated proteins (using biotin). PPBD was not included in the control cells. The scale bars represent 10 µm. (C) Intensity profile of line in panel 1B. (D) Mascot score of proteins detected in the experimental sample (with PPBD) versus the control sample (without PPBD). The top 22 proteins are highlighted in red. The already-known speckle resident’s name is included. (E) Localization analysis of the polyP interactome using the g:Profiler database between cytoplasm and nucleoplasm. (F) Nuclear localization analysis of the PPP (polyP Proximity Proteome). (G) Functional analysis of the PPP using the ShinyGO 0.80 database. The –log_10_FDR (False Discovery Rate) of each comparison and the number of hits is represented. (H) Main domains present in the PPP along with the identifier in the Interpro database, the number of proteins in the list containing it, its percentage, and the FDR value. (I) Polyphosphate colocalizes with nuclear speckles. Immunolocalization of SRRM2 (speckle), PML (PML bodies), coilin (Cajal bodies), fibrillin (nucleolus) and H3K9me3 (heterochromatin). The scale bars represent 10 μm. (J) Pearson correlation coefficient of the colocalization between polyP and the different condensates in I (K) Venn diagram comparing PPP and the SRRM2 proximity proteome both obtained using BAR. The graph shows the colocalization Pearson correlation coefficient. At least 40 cells from three different experiments were included in the analysis.

### Polyphosphate interacts with the core NS element SRRM2

To identify the specific NS element interacting with polyP, we hypothesized that PASK domains might play a role, as these regions have been implicated poly-phosphorylation (36). Using a custom Python script, we searched for PASK domains within our polyP proximal proteome (PPP). Remarkably, only three proteins exhibited a clear PASK domain, with SRRM2 showing the highest score value and followed by SON (Fig. 2A). Notably, both proteins, within their PASK region, contain distinct lysine-rich stretches described as important regions for polyP interaction (38). In SRRM2, this domain overlaps with a highly disordered and highly polar sequence (Fig. S2A), making it an excellent candidate for interaction with a polyanionic molecule such as polyP. Consistent with our algorithm’s prediction, EMSA (Electrophoretic Mobility Shift Assay) analysis demonstrates that polyP interacts *in vitro* with the SRRM2 PASK region (Fig. 2B). Polyphosphate also binds SON PASK region although showing a weaker interaction (Fig. 2B). To further investigate the polyP-SRRM2 *in vitro* interaction, we characterized the polyP binding properties of SRRM2 domains. SRRM2 contains two disordered and polar Arg-Ser (RS) repeat regions essential for maintaining NS integrity. However, these regions do not interact with polyP (Fig. 2B), highlighting the specific binding affinity between polyP and the SRRM2_PASK_ region. Notably, the SRMM2_PASK_/polyP interaction is markedly reduced when the 7xLys stretch is mutated to 7xAla (Fig. 2B) suggesting a prominent role for the 7xLys sequence (Fig 2B). The interaction is independent of the polyP size except for the shortest polyP 3-mers, which show negligible binding (Fig. 2C). Finally, while RNA binds both the PASK and RS1 regions, as previously described (49, 51) and (Fig. S2B), polyP exhibits specific and selective binding to the PASK region, highlighting its preference for polyP over RNA.

**Figure 2.**
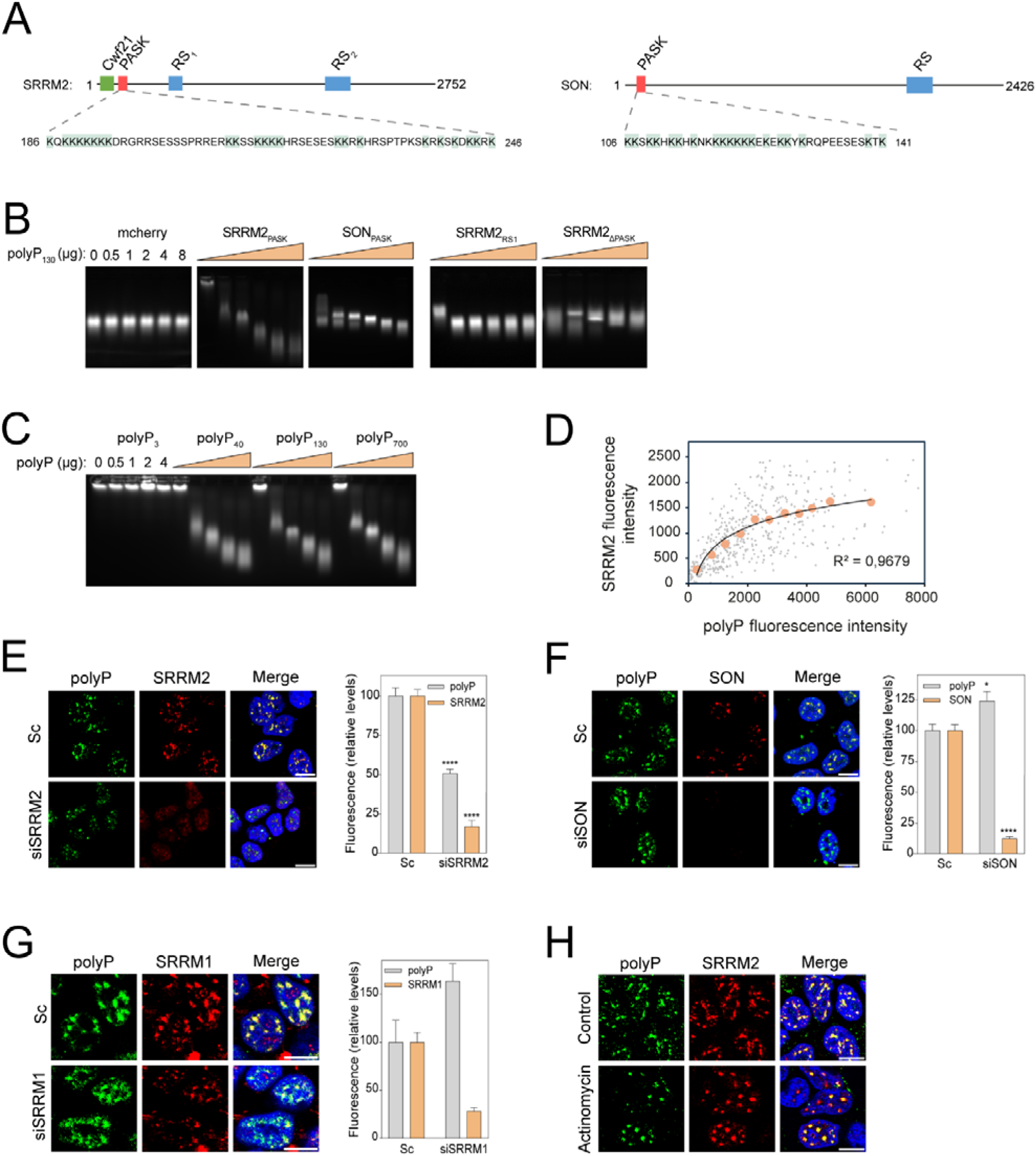
Polyphosphate interacts with the PASK domain of SRMM2. (A) SRRM2 and SON domains map highlighting the K-rich sequence in the PASK domain. (B) Polyphosphate interacts with PASK. EMSA analysis of growing amounts of polyP with the SRRM2 PASK and RS recombinant domains. The mobility of the protein was observed by the mCherry fluorescence. (C) EMSA including PASK and a growing amount of different polyP sizes. (D) Polyphosphate and SRRM2 amounts correlate in individual cells. SRRM2 and polyP were immunostained and quantified in a total of 500 cells. Cells were grouped according to polyP intensity in groups of 500 intensity units. Mean± SEM of polyP and SRRM2 of those groups is presented and adjusted to a logarithmic function (orange dots). (E, F, and G) NS localization of polyP depends on NS proteins. Polyphosphate immunolocalization in cells where SRRM2, SON or SRRM1 have been targeted with specific siRNA. After 48 h, polyP and the different NS proteins were detected and quantified. A minimum of 500 cells per condition were analyzed. Mean intensity ± SEM of at least fifteen pictures from three independent experiments. (H) Inhibition of transcription rearranges polyP in the speckle. Cells were treated with 10 μg/μL of actinomycin. After 10 h, SRRM2 and polyP were detected as in E. *p < 0.05, **p < 0.01; ***p < 0.001; ****p < 0.0001. Scale bars represent 10 μm.

To check the *in vivo* relationship between polyP and SRRM2 predicted by the EMSA *in vitro* experiments, we performed immunolocalization studies at the single-cell level that reveal a correlation between SRRM2 and polyP abundance, with polyP acting as the limiting factor (Fig. 2D). Additionally, the depletion of SRRM2 dramatically reduces the amount of polyP in the NS without affecting other NS elements (Fig. 2E and S2C). Interestingly, the depletion of SON and SRRM1 triggers and increase in SRRM2 (Fig. S2C), resulting in a higher polyP content in the NS (2F and 2G) supporting the reduced interaction found in the *in vitro* experiments. Finally, SRRM2 and polyP behave similarly and change their shape coordinately to a round and sharply punctuated pattern when cells are treated with actinomycin, a transcription inhibitor that induces such a variation in the shape of the NS (Fig. 2H and (52)). All the above results confirm a differential interaction between polyP and NS elements, highlighting the PASK region of SRRM2 as the main recruiting factor for polyP in the NS.

Considering the prominent role of NS in mRNA splicing processes, it is timely to investigate the possible role of polyP in splicing.

### Polyphosphate regulates splicing

NS have been involved in several functions, but those related to RNA splicing are prominent (5) indicating that polyP might be regulating this cellular process. Coherently, splicing is the first GO function group obtained in the PPP presented above (Fig. 1G). As a first approach to assess the role of polyP in cellular splicing, we evaluated intron removal using a luciferase reconstitution assay ((50) and Fig. 3A). As expected, splicing efficiency is significantly reduced in the presence of the pre-mRNA splicing inhibitor isoginkgetin (53), validating the system (Fig. 3B). Interestingly, degradation of polyP via overexpression of either Ppx1 or NUDT3 results in a splicing defect very similar to the depletion of SRRM2 (Fig. 3B). This result prompted us to analyze the differential alternative splicing events (DASEs) by RNA-seq in cells overexpressing Ppx1 (which reduces polyP levels, leading to SRRM2 delocalization) and in SRRM2 downregulated cells. Regarding their respective DASEs (Supplemental Table 2 and 3), the two genetic models produce a similar proportion of differentially spliced events and exon skipping represents the most frequent event (Fig. 3C and S3A). The RT-PCR analysis of events with a high ΔPSI value validates the RNA-seq analysis (Fig. 3D).

**Figure 3.**
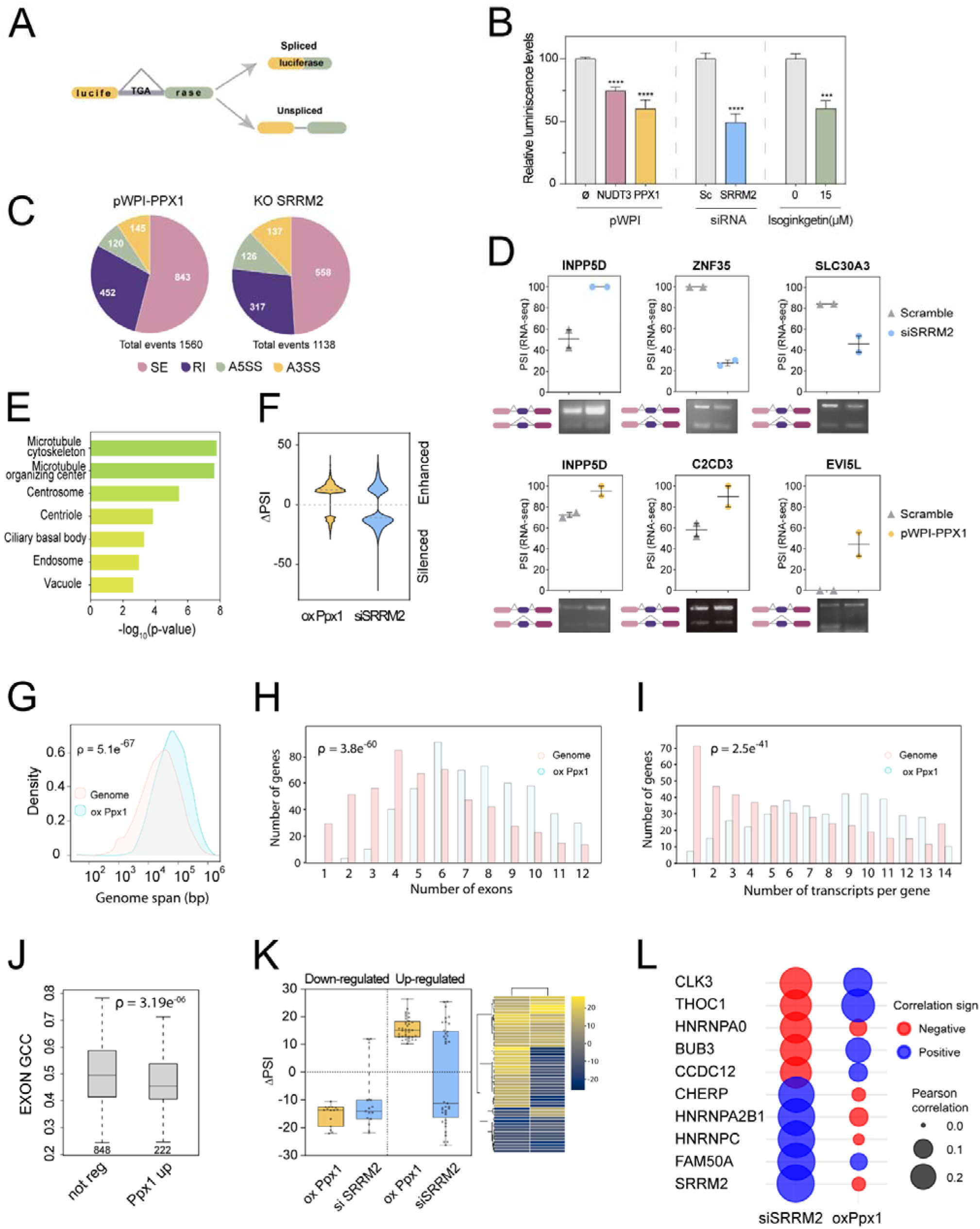
Polyphosphate regulates splicing. (A) Diagram for the splicing efficiency experiment. (B) Splicing is impaired in cells with reduced levels of polyP. HEK293T cells were transfected with empty pWPI, pWPI-NUDT3, or pWPI-Ppx1 together with the pCMV-LUC2CP vector. After 72 h, luciferin was added to each well and luminescence was measured. SRRM2 silencing or isoginkgetin treatment were used as positive controls. The graph shows the mean ± SEM of five independent experiments. (C) siSRRM2 cells and polyP-depleted cells have similar differentially spliced events patterns. Diagram showing the percentage of differential alternative splicing events in the RNAseq datasets of cells overexpressing Ppx1 and in cells where SRRM2 has been silenced. SE is for exon skipping, RI for intron retention, A5SS for alternative 5’ splice-site and A3SS for alternative 3’ splice-site (D) RNAseq validation. Randomly selected events in the two RNAseq were analyzed by RT-PCR. Representative images of the agarose gel analysis of the fragments with the PSI obtained from gel quantification along with a map of the event are provided. The graphs are the PSI values obtained in the RNAseq analysis for these genes. (E) Depleting polyP mainly affects the exon skipping of genes involved in microtubules. GO subcellular function of exon skipping events in cells with reduced levels of polyP by overexpressing Ppx1. (F) Analysis of exon skipping events in SRRM2 silenced and polyP depletion datasets. Up-regulated (>10 ΔPSI) is for exon inclusion and down-regulated (<-10 ΔPSI) is for exon exclusion in the noted datasets. (G) Large genes are mainly affected by polyP depletion. Genome span refers to specific regions covered by a particular gene. ShyniGO analysis of polyP depleted (ox Ppx1) dataset vs standard genome. (H) Transcripts with numerous exons are enriched in polyP-depleted cells dataset. ShyniGO analysis as in G. (I) Transcripts with a high number of isoforms are enriched in polyP depleted cells dataset. ShyniGO analysis as in G. (J) The GC content of affected exons in the polyP depleted cells dataset. Up-regulated and down-regulated exons with ΔPSI>=15 were compared against a randomly selected subset of un-regulated exons using the MATT pipeline (77). Ppx1 up is for inclusion events in F and Ppx1 down is for exclusion events in F. (K) Analysis of coregulated events in siSRRM2 and polyP depleted cells datasets. Shared events in Fig. S4E were considered. Ppx1 events were separated into exclusion (down-regulated) and inclusion (up-regulated) events. For every Ppx1 event, the corresponding siSRRM2 is represented (left panel). Heatmap of the above analysis (right panel). (L) Correlation analysis between ΔPSI of regulated exons in siSRMM2 or polyP-depleted cells and a siRNA array of >250 splicing-related proteins. Only top correlating factors are shown.

The molecular function GO analysis of the subset of exon skipping events reveals that the polyP-dependent splicing targets are related to centrosome and microtubule biology (Fig. 3E). Interestingly, the same enrichment in centrosome elements is observed when SRRM2 is depleted (Fig. S3B and consistent with (51) showing, again, a close functional relation between polyP and SRRM2. Note that the SRRM2 and polyP functional relationship is supported by the fact that 25% of the affected genes are shared in polyP and SRRM2 depleted cells (Fig. S3C).

From a molecular perspective, polyP depletion mainly increases exon-inclusion, while SRRM2 depletion predominantly promotes exon-skipping (Fig. 3F). The bias toward exon-inclusion in SRRM2 physiological role, points to a possible role of polyP in releasing splicing factors from NS.

The analysis of specific polyP-regulated events shows that the included exons predominantly belong to large genes (Fig. 3G) with a high number of exons (Fig. 3H) present in long transcripts (Fig. S3C) and with many isoforms (Fig. 3I). Additionally, both the exons and the adjacent introns (upstream and downstream) exhibit low GC content (Fig. 3J and S3D). Considering that the pre-mRNAs of large genes are, mainly, spliced far from NS (21), the results indicate that polyP depletion triggers the pre-mRNA processing of long transcripts distant from NS. Comparing polyP-depleted and SRRM2 downregulated exon splicing datasets shows a 10% overlap of splicing events, which represents the putative coregulated events both by polyP and SRRM2 (Fig. S3E). The analysis of the coregulated events evidences a positive correlation in the skipped exons across both datasets (Fig. 3K) which is compatible with the notion that polyP is needed to maintain SRRM2 in the NS. On the contrary, half of the up-regulated exons upon polyP depletion show increased skipping upon SRRM2 silencing, suggesting that the SRRM2 activity in processing certain types of transcripts relies on polyP degradation (Fig. 3K). These findings, along with those presented in previous Figs., support the hypothesis that polyP regulates the dynamic localization of SRRM2, modulating the processing of short transcripts within NS while long transcripts are processed farther away.

Splicing regulates the coordinated action of more than 300 proteins, including core spliceosome components, splicing factors and splicing factor kinases among others (54). Rogalska et al. coupled the KD of more than 250 splicing regulatory proteins with RNAseq, allowing the identification of splicing events regulated by specific factors and highlighting functional relationships between factors in an unbiased approach (54). To identify functional relationships between polyP and other splicing regulatory factors, the ΔPSI values obtained in the Rogalska collection were confronted with the siSRRM2 dataset finding that SRRM2 displays the highest positive Pearson correlation (Fig. 3L), further validating the comparison of the datasets. Notably, the top negative Pearson correlation corresponds to CLK3, suggesting that this protein may act in opposition to SRRM2. Remarkably, when comparing our Ppx1 overexpression dataset to the above collection, CLK3 again emerges again as one of the most correlated events (Fig. 3L), pointing to a potential genetic relationship between polyP, SRRM2, and CLKs.

### Polyphosphate restricts CLKs activity

CLK kinases regulate the intranuclear distribution of SR proteins influencing Pre-mRNA splicing (55). Accordingly, the overexpression of CLK1 triggers the disassembly of splicing speckles (20). We first demonstrated that recombinant CLK3 is active against a short peptide containing four RS repeats (56) (Fig. S4A) and that retains the ability to be regulated by temperature described for the CLK1 kinase (Fig. 4A and 4B, (57)). This result suggests that both proteins might be regulated through similar mechanisms. Importantly, CLK3 activity is efficiently inhibited by both short (14-mer) and long (130-mer) polyP molecules (Fig. 4C and S4B) and this inhibition also occurs against the most well-characterized substrate of CLK1: the SRSF1 protein (Fig. 4E). Interestingly we demonstrated for the first time that the RS-enriched region of SRRM2 (the RS_1_ region, Fig 2A) is an *in vitro* substrate of CLK3 kinase and that this phosphorylation is again inhibited by the presence of polyp in a dose dependent manner (Fig. 4D, and S4C). Lastly, the same results are obtained when CLK3 is produced in insect cells, indicating the absence of any posttranslational requirement for CLK3 polyP inactivation (Fig. S4D). It is worth noting that polyP does not inhibit CDK16 (Fig. 4F), a kinase with structural similarities to CLK3 that also belongs to the CMGC kinase family (58, 59) ruling out the possibility of non-specific inhibition in the *in vitro* enzymatic assay and suggesting that polyP inhibition of CLK3 is specific. Besides, the fact that the increase of ATP (substrate) in the reaction does not reduce the inhibitory effect of polyP on CLK3, suggests a non-competitive inhibition (Fig. 4G). Furthermore, the reduction of polyP levels results in notable *in vivo* phosphorylation of various SRSF proteins (Fig. 4H), well-known targets of CLK kinases (56). On the contrary, incrementing the levels of polyP by expressing the bacterial polyP synthase PPK (60) inhibits the CLK1 activity even when overproduced (Fig. 4I). All the results presented above presents polyP as a potential physiological inhibitor of CLK kinases.

**Figure 4.**
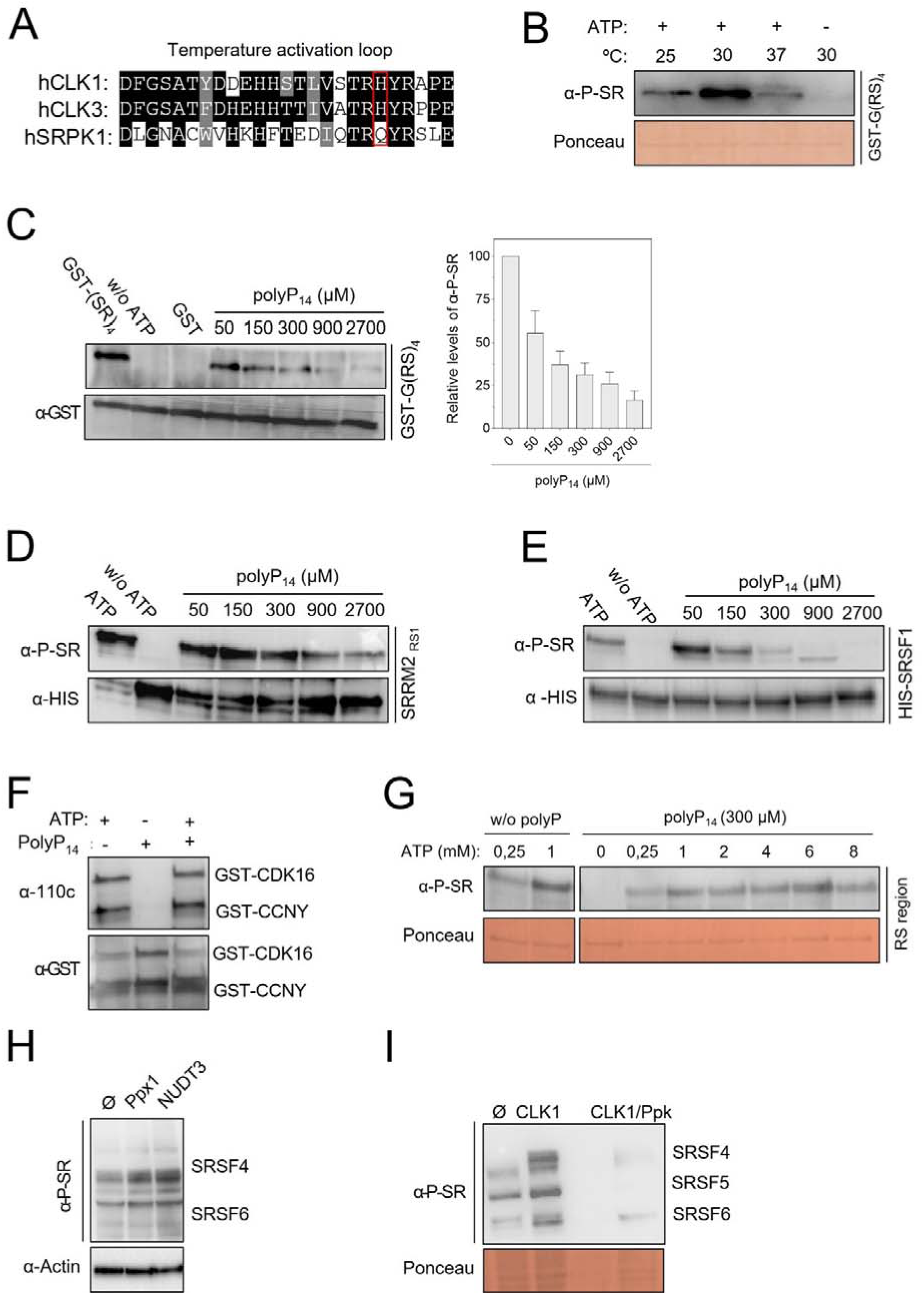
Polyphosphate inhibits CLKs activity *in vitro* and *in vivo*. (A) CLK3 bears a temperature activation loop. Sequence alignment of CLK3 with the temperature activation loop of CLK1 and SRPK1. Marked in red the His essential for the temperature regulation. (B) Temperature regulation of the CLK3 kinase activity on recombinant SR repeat peptide. Representative western blot using specific anti-phospho SR antibodies. Recombinant CLK3 was assayed at the noted temperatures using as substrate a peptide containing 4 repetitions of the RS sequence. (C, D and E). CLK3 is inhibited by polyP. Representative western blots as in B of CLK3 activity analyzed in the presence of growing amounts of polyP_14mer_ using as substrate a four repetition of the RS sequence, the RS region of SRRM2 or SRSF1. (F) Polyphosphate does not inhibit CDK16 kinase activity. Kinase assay using recombinant CDK16 and its activator cyclin CCNY. CDK16 autophosphorylation and phosphorylation on CCNY were detected using a specific anti-thiophosphate ester antibody. (G) ATP does not compete with polyP for the CLK. Western blot representative. The same kinase assay as in D but adding increasing amount of ATP. (H) Polyphosphate i*n vivo* inhibits SR proteins phosphorylation. Polyphosphate was depleted by overexpressing Ppx1 and NUDT3, a western blot of cell extracts using specific anti-phospho-SR antibodies. (I) CLK1 activity is inhibited by polyP *in vivo*. Representative western blot as in H. In cells overexpressing PPK for 24 h and controls, CLK was overexpressed for 24 h.

### NS integrity is controlled by polyP/CLK axis

CLK phosphorylation on SR proteins has been shown to produce the NS disassembly (16). Because of our later results, we predict that the modulation of the nuclear polyP levels should affect the NS integrity through the activation of CLK proteins, and this is the case indeed. The overexpression of both Ppx1 or NUDT3 significatively reduce the polyP and concomitantly the NS (Fig. 5A, 5B and S5A), without affecting SRRM2 protein levels (Fig. S5B). The same effect appears in a dose-dependent manner when NUDT3 overexpression is regulated (Fig. 5C). Importantly, the effect on the NS by polyP depletion is abolished in the presence of T3, a specific inhibitor of CLKs (61) (Fig. 5D). Also predicted by our model, incrementing polyP levels both by silencing NUDT3 or by overexpressing Ppk1 leads to an increase of NS (Fig. 5E, S5C and S5D). Lastly, the increase in nuclear polyP makes cells resistant to CLK1 overexpression-induced NS disintegration (Fig. 5F). Collectively, these experiments demonstrate that polyP regulates CLK activity and thereby orchestrates the mobilization of splicing factors from NS.

**Figure 5.**
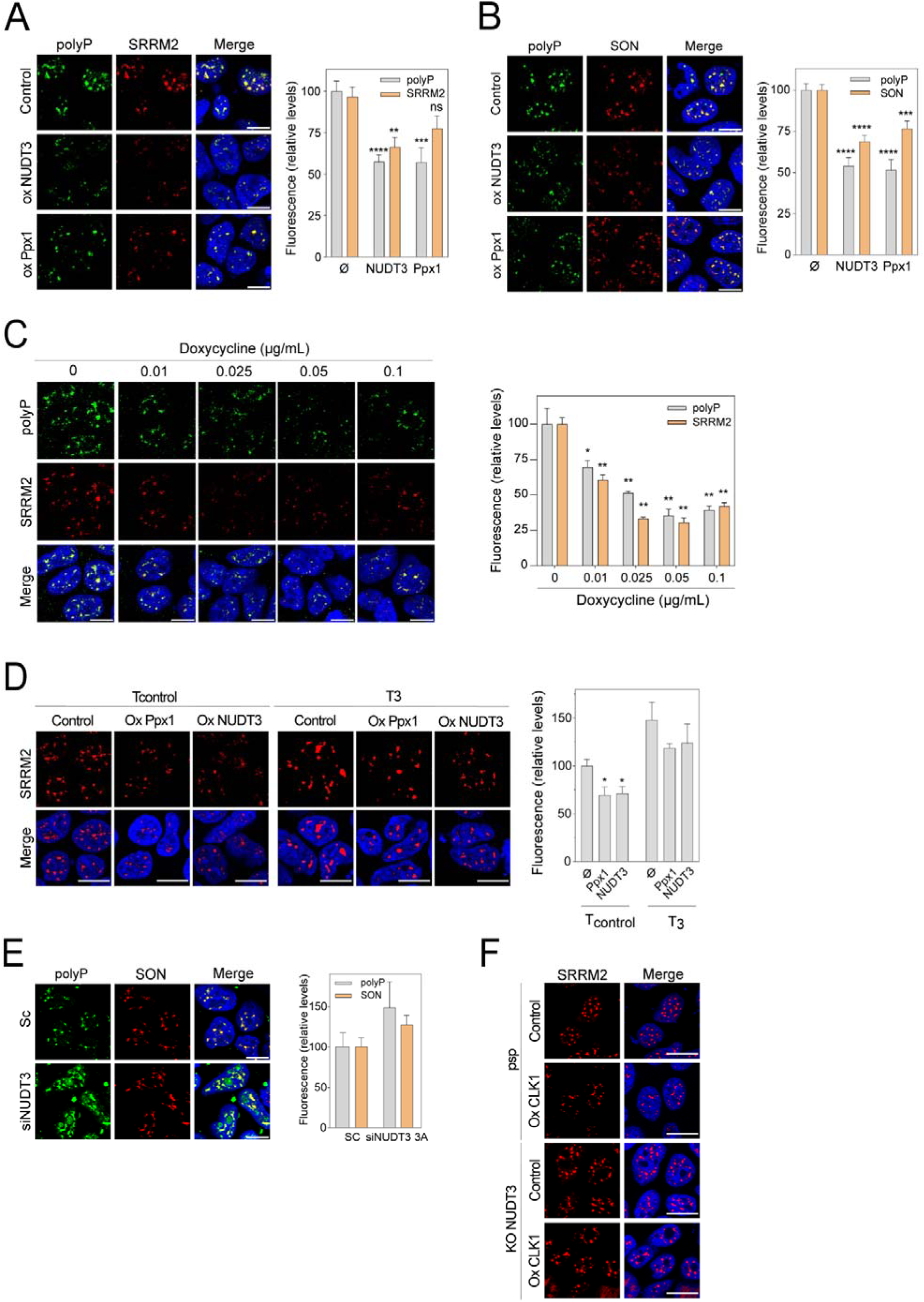
Polyphosphate regulates NS morphology. (A, B) Polyphosphate constitutive depletion delocalizes SRRM2 and SON. Cells were transfected with empty pWPI, pWPI-NUDT3 (human endopolyPase), or pWPI-Ppx1 (exopolyPase). After 48 h polyP, SRRM2 (A) and SON (B) were quantified by immunofluorescence. A minimum of 500 cells per condition were analyzed. Mean intensity ± SEM of at least fifteen pictures from three independent experiments. (C) SRRM2 delocalizes upon polyP depletion in a dose-dependent manner. Cells were transfected with empty pRetroX or pRetroX-hNUDT3. NUDT3 gene expression is controlled by adding increasing amounts of doxycycline. After 48 h, polyP and SRRM2 were quantified by immunofluorescence. Quantification of mean intensity ± SEM of five pictures from one experiment. (D) Polyphosphate regulation of NS is CLKs dependent. Polyphosphate depleted cells by the overexpression of Ppx1 or NUDT3 were treated for 24 h with T3, a CLKs inhibitor, or T-an inactive analog. SRRM2 was quantified by immunofluorescence. A minimum of 500 cells per condition were analyzed. Mean intensity ± SEM of at least five pictures from two experiment. (E) Polyphosphate increase results in SRRM2 accumulation. Cells were transfected with scrambled or NUDT3-targeting siRNA. After 72 h, polyP, SRRM2 was quantified by immunostaining. A minimum of 500 cells per condition were analyzed. Mean intensity ± SEM of at least ten pictures of one experiment. (F) Polyphosphate induced NS increment is CLK1 dependent. CLK1 was overexpressed in NUDT3 knock-down cells. SRRM2 was quantified by immunofluorescence. A representative image of four independent experiments. *p < 0.05; **p < 0.01; ***p < 0.001; ****p < 0.0001. The scale bars represent 10 μm.

### Model and physiological relevance

We propose a model in which polyP associated with SRRM2 works as a sort of shield inhibiting the CLKs, thus preventing the CLK-mediated phosphorylation of SR domains and, in turn, keeping the NS assembled and visible (Fig. 6A). It is known that oxidative stress reduces polyP levels (43), making it a physiological condition to test the proposed model. The addition of menadione, besides reducing polyP levels as previously described, significantly promotes NS disassembly (Fig. 6B). Interestingly, polyP depletion precedes SRRM2 delocalization, suggesting a time-dependent regulatory mechanism. NS disorganization under oxidative stress is entirely dependent on NUDT3 and subsequent CLK activities, as demonstrated by two key observations. First, NUDT3 KO cells, which are unable to degrade polyP, do not exhibit NS mobilization under stress (Fig. 6C). Second, even when polyP is degraded, NS disorganization does not occur if CLK activity is inhibited (Fig. 6D).

**Figure 6.**
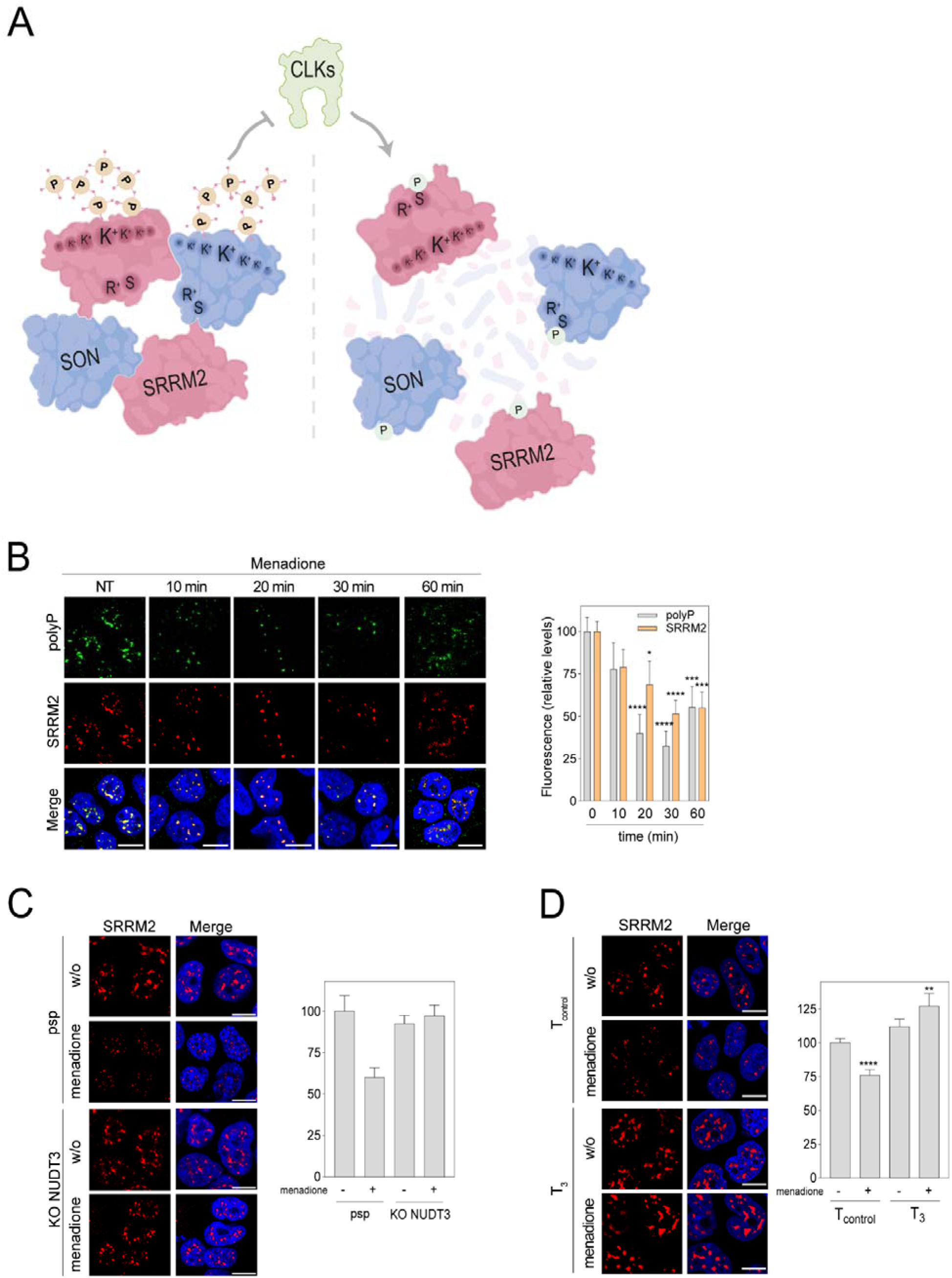
Working model. Polyphosphate binds to the PASK domains of nuclear speckles (NS) proteins SRRM2 inhibiting CLK kinase activity and maintaining NS in a tight configuration (left). When polyP levels are reduced, CLK kinases regain their activity and phosphorylate SR proteins, leading to the NS disassembly and the release of splicing factors to the nucleoplasm (right). (B) Polyphosphate reduction upon oxidative stress is accompanied by a reduction in SRRM2. Cells were treated with 15 μM menadione for the noted times, immediately fixed and immunostaining of SRRM2 and polyP was performed. Mean intensity ± SEM of at least fifteen pictures from three independent experiments. (C) Oxidative stress induced SRRM2 reduction is NUDT3 dependent. NUDT3 knock-down cells were treated with 15 μM menadione for 30 min. SRRM2 was quantified by immunostaining. Mean intensity ± SEM of five pictures (at least 500 cells condition were analyzed) from one experiment. (D) Cells were treated with 15 μM menadione for 30 min. in the presence of the inhibitor T3 or its inactive analog T-. SRRM2 was quantified by immunostaining. Mean intensity ± SEM of at least fifteen pictures from three independent experiments. *p < 0.05; **p < 0.01; ***p < 0.001; ****p < 0.0001. The scale bars represent 10 μm.

In line with our model, under specific stress conditions, the degradation of polyP is essential for activating CLK proteins. These proteins phosphorylate NS elements, enabling their mobilization to facilitate the splicing of stress response RNAs outside of speckles.

## DISCUSSION

### The polyP proximal proteome: a novel approach to understanding human polyP functions

The polyP role in human cells has been difficult to define, largely due to the challenges in characterizing its metabolic machinery and in isolating it from cells and tissues. To clarify its function, we aimed to identify the proteins interacting with polyP. Three critical issues must be considered when pursuing this strategy: First, polyP is highly negatively charged, which causes it to interact non-specifically with basic proteins, making co-immunoprecipitation (CoIP)-based approaches problematic (62). Second, polyP has a high turnover rate (39) making difficult the detection in approaches requiring stable interactions. Third, polyP lacks a defined spatial structure, and despite efforts, no sufficiently specific antibodies have been produced (63). Proximity labelling strategies, however, can overcome these challenges by rapidly tagging transient interactions and enabling analysis in intact and metabolically arrested cells. Using the high specificity of the PPBD peptide in binding to polyP (42, 43), we adapted the BAR technique to generate a reliable polyP Proximal Proteome (PPP). Several results in this study (Fig. 1 and Fig. S1) validate the approach, demonstrating for the first time that BAR can be applied to detect the proximal proteome of metabolites if an appropriate sensor for the specific metabolite is available.

The first major finding from the PPP is that polyP localizes in several nuclear membrane-less organelles. Here, we focused on the polyP role in regulating splicing within NS; however, our findings open new avenues for investigating the polyP functions in other locations, particularly in the nucleolus, where polyP has been previously reported (60, 64). Additionally, we have found polyP in chromatin, Cajal bodies, and the nuclear membrane (Fig. 1F), suggesting that polyP may play roles yet to be explored in these structures. The widespread presence across various nuclear condensates points to polyP as a potential common structural or regulatory factor in these highly dynamic cellular compartments.

### NS from a membrane-less condensate to a complex structure

Initially, nuclear condensates were considered disordered accumulations of molecules gathered by phase separation (3). However, focusing on NS, alternative models have emerged, including a complex conformation of different elements mediated by specific binding interactions, highlighting the role of electrostatic forces supported by polyanions such as RNA (including long non-coding RNAs) and DNA (3, 65). The nucleation of NS via mechanisms like the spatial clustering of active genes reinforces the notion of the high complexity of these structures (66). Moreover, modern high-resolution microscopy reveals intricate layers within NS (67), as well as coordinated protein subdomains (13). Undoubtedly, future research will uncover additional components in the NS network, further expanding our understanding of this organelle.

In this context, we propose polyP as a novel component of NS. Functionally, polyP stands out as the only known factor whose depletion leads to NS disassembly. While we propose that polyP interacts with SRRM2 through its PASK domain, other proteins may also recruit polyP to the NS. Indeed, two more proteins in our PPP (SON and THOC2) contain robust PASK domains that could mediate polyP recruitment. Additionally, DYRK1A, a protein found in NS (38), contains a poly-His stretch that might also serve as an anchoring site for polyP in NS. Whether polyP is synthesized near NS or transported to NS remains an open question; addressing it will be crucial for a more complete understanding of NS dynamics.

### Polyphosphate as a new splicing regulator

The nucleus can be divided into two transcriptional hot zones: one located within NS predominantly found in the central region of the nucleus and another in the nuclear periphery (68). This highlights the need for a precise distribution of splicing factors to meet the cell transcriptional demands. In this sense, it is well established that splicing factors reside in NS for approximately 50 sec, indicating continuous shuttling between NS and the nucleoplasm (69). Notably, the splicing of large genes generally occurs outside the NS (21), and our data show that polyP depletion leads to increased exon inclusion in those large genes. These findings suggest a complex regulatory mechanism governing the distribution of splicing factors throughout the nucleus, further supporting polyP’s role as a key regulator in the mobilization of splicing factors.

### A model for polyP-mediated splicing control

SR proteins contain RS repeat domains essential for maintaining the NS structure (70, 71). These domains are targets for phosphorylation contributing to NS disassembly (16). CLKs, along with SRPKs and DYRKs, are the kinases responsible for regulating the phosphorylation of SR proteins (72). Interestingly, our study identified CLK3, the least known member of the Ser/Thr kinase CLK family (73), as a target for polyP inhibition.

It is known that polyP inhibits DYRK1A by interacting with its poly-His stretch (38). Since CLK3 lacks clear poly-His or poly-Lys stretches, the exact mechanism by which polyP inhibits CLK3 kinase activity remains unclear and, therefore, we cannot rule out the possibility that polyP inhibitory effect could be extended to other CLK family members. Given that CLKs are more promiscuous in substrate selection compared to SRPKs (74), and considering that CLKs localization is compatible with NS (kinome_atlas http://cellimagelibrary.org/pages/kinome_atlas), a mechanism inhibiting the CLKs gains relevance in the NS control. Therefore, we propose a model in which polyP modulates splicing by regulating CLK kinase activity (Fig. 6A). When polyP is present above a critical level (e.g., bound to PASK domains in SRRM2 and SON proteins), CLK kinase activity is restrained, preventing the phosphorylation and disassembly of SR proteins. However, when splicing activity needs to be adjusted (such as for the splicing of large genes), a reduction in polyP levels makes SR proteins more vulnerable to CLK action, leading to their mobilization from the NS and, eventually, to NS disassembly.

One well-documented instance of polyP level regulation occurs during oxidative stress, where nuclear polyP is degraded by the activation of the endopolyphosphatase NUDT3 (43). According to our model, polyP depletion leads to the displacement of SRRM2 enabling cells to adapt and respond. A similar polyP-regulated mechanism could occur during the cell cycle. It is known that NS dissolve in early mitosis through the activity of CLK1 and DIRK3, and reform later in the cell cycle. (19, 75, 76). In both cases, we speculate that the degradation and resynthesis of polyP could be the key factor triggering the assembly/disassembly cycles of NS. The fact that polyP is cyclically regulated in lower eukaryotes (28) helps support this speculation.

In summary, we propose a model in which the highly dynamic nature of NS is regulated by polyP through its inhibition of CLK activity. Our model suggests that by modulating splicing factors localization, polyP may exert control over the wide array of cellular functions in which it is involved. This new role of polyP opens new avenues for understanding how cells fine-tune gene expression in response to various physiological conditions, positioning polyP as a critical player in nuclear organization and RNA processing.

### Limitations of the study

It must be noted that BAR labels cellular components within a radius of influence (usually 20 nm). Then not all the already described NS elements have been found.

The work has been performed in HEK293T human model cells. Derived from the fact of polyP amount variations according to cell type (60), polyP function could vary importantly among cell types and tissues.

## DATA AVAILABILITY

All data will be available in a subject-specific repository. Accession numbers and identifiers will be provided.

## SUPPLEMENTARY DATA

Supplementary Data are available online.

## AUTHOR CONTRIBUTIONS

Blanca Lázaro: Investigation, formal analysis. Francisco J. Tadeo-Masa: Investigation, formal análisis. Andrea Rodriguez: Investigation. Lucia Ayuso: Investigation. Joan M Martínez-Láinez: Investigation. Eva Quandt: Investigation. Maribel Bernard: Investigation. Filipy Borghi Investigation. Adolfo Saiardi: Methodology, validation. Jonàs Juan-Mateu: Formal analysis, data curation, validation. Javier Jiménez: Conceptualization funding adquisition writing original draft review and editing. Josep Clotet: Conceptualization, funding adquisition, supervision, validation writing and original draft review and editing. Samuel Bru: Conceptualization, investigation, supervision, validation writing and original draft review and editing

## ACKNOWLEDGEMENTS

The proteomic analysis was performed in the Proteomics Unit of Complutense University of Madrid. We thank Marta Pérez, Meritxell Martínez and Paula Rodríguez for their technical support. This work was supported by and is part of the I+D+i grant ref: PID2021-127302NB-I00 to JC and JJ by the Spanish Ministerio de Ciencia e Innovación (MCIN).

## FUNDING

This work was supported by the Ministerio de Ciencia e Innovación. Spanish Government [PID2021-127302NB-I00 to JC and JJ] Funding for open access charge: Ministerio de Ciencia e Innovación.

## CONFLICT OF INTEREST

The authors declare no conflict of interest.

## Notes

### Competing Interest Statement

The authors have declared no competing interest.

