## Supplementary material for "Nuclear Speckle Dynamics are Controlled by Polyphosphate Inhibition of CLK Proteins": supplemental inf.docx

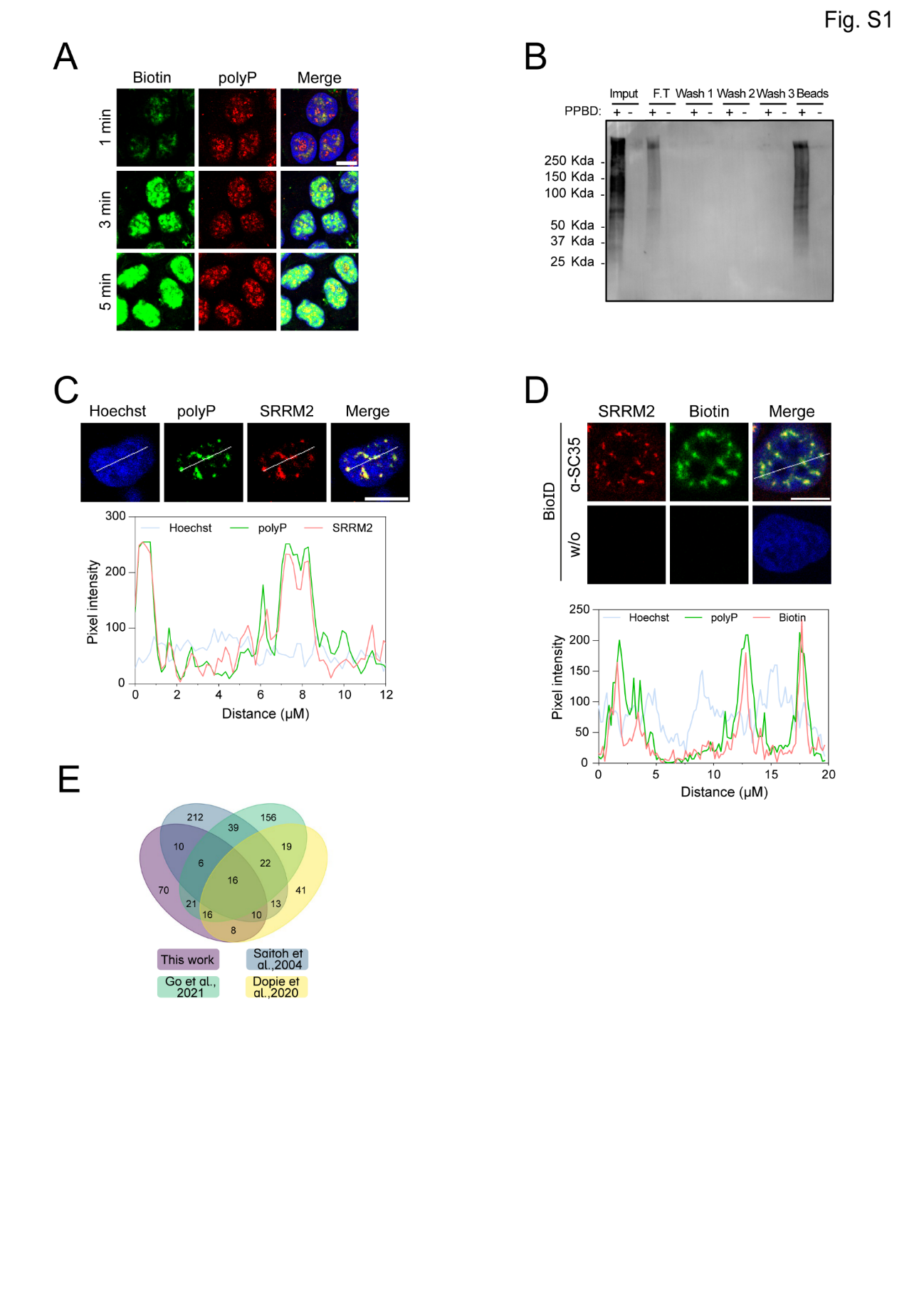
**Figure S1.** (A) Biotinylating reaction time set up. Cells were fixed and treated as described in the material and methods. The biotinylating reaction was stopped at the indicated times. polyP and biotin were analyzed by immunofluorescence. (B) Western blot of the BAR different steps. Samples of the indicated steps in the biotinylating process were loaded in an SDS-PAGE gel, blotted and detected using a specific α-biotin primary antibody coupled to a secondary α-mouse antibody. The biotinylating process without PPBD was used as a control. (C) Confocal images of immunolocalized SRRM2 and polyP (Upper). SRRM2, polyP and Hoechst intensity profiles from the line in the upper panel (Lower). (D) Confocal images of cells labeled with α-SRRM2 and α-biotin (Upper). Cells without the α-SRRM2 were used as a control. Biotin, SRRM2, and Hoechst intensity profiles from the line in the upper panel (Lower). (E) Venn diagram of the SRRM2 interactome in this work and three other interactomes published for SRRM2 (1, 2). Scale bars represent 10 µm.

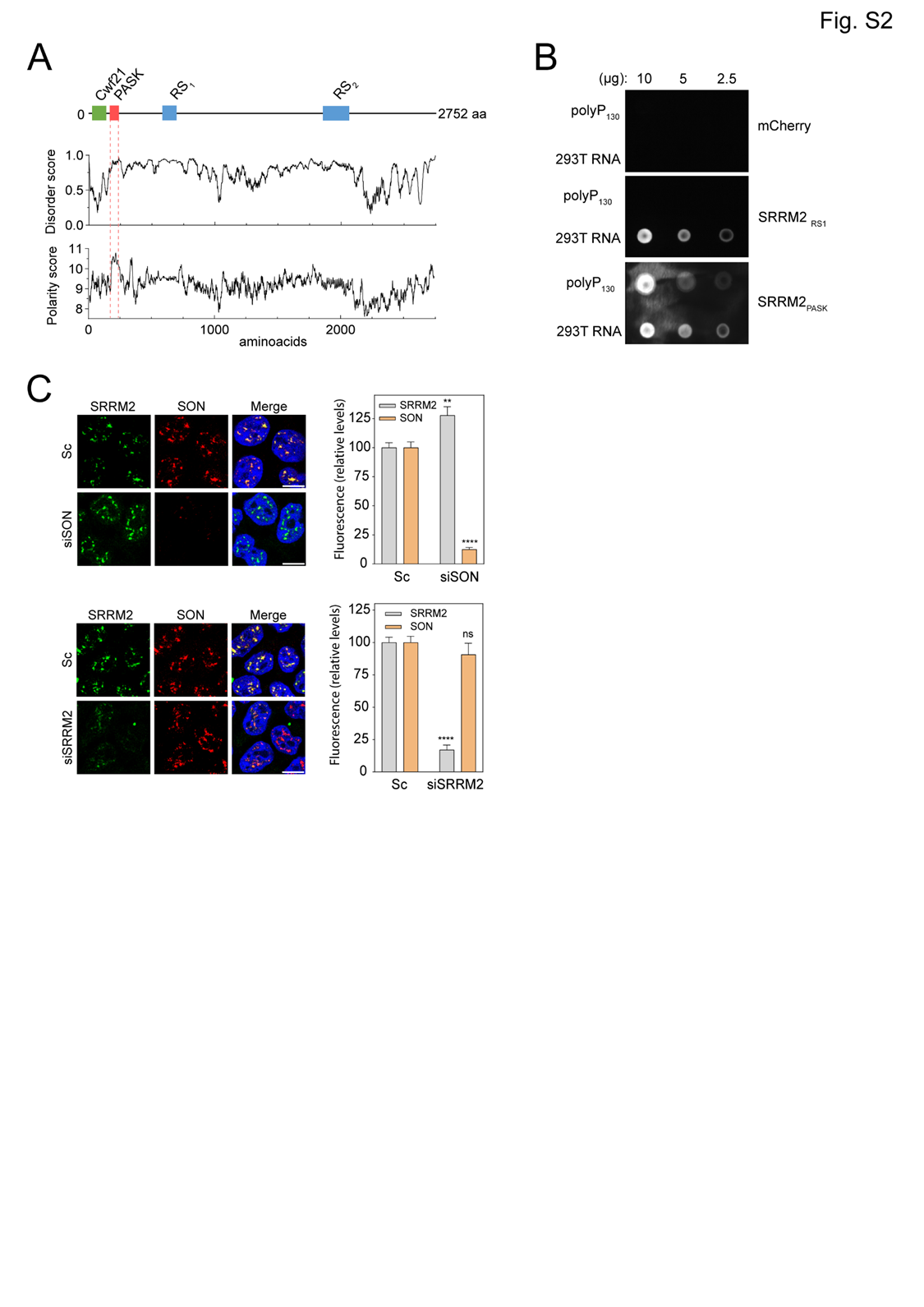

**Figure S2.** (A) SRRM2 main structural domains emerging from the disorder and polarity profiles of SRRM2 sequence. The Cwf21 domain (IPR013170), the PASK domain and the RS_1_ and RS_2_ repeat regions (3) are depicted. (B) Dot blot showing the binding of PASK and RS domains (mCherry-tagged) to different amounts of polyP_130_ and RNA. mCherry was included as a control. (C) SRRM2 and SON immunolocalization. Cells were transfected either with scramble, siSRMM2 or siSON. SRRM2 and SON localization was analyzed after 48 h. A minimum of 500 cells per condition were analyzed. Mean intensity ± SEM of fifteen pictures of three independent experiments is shown. **p < 0.01; ****p < 0.0001.

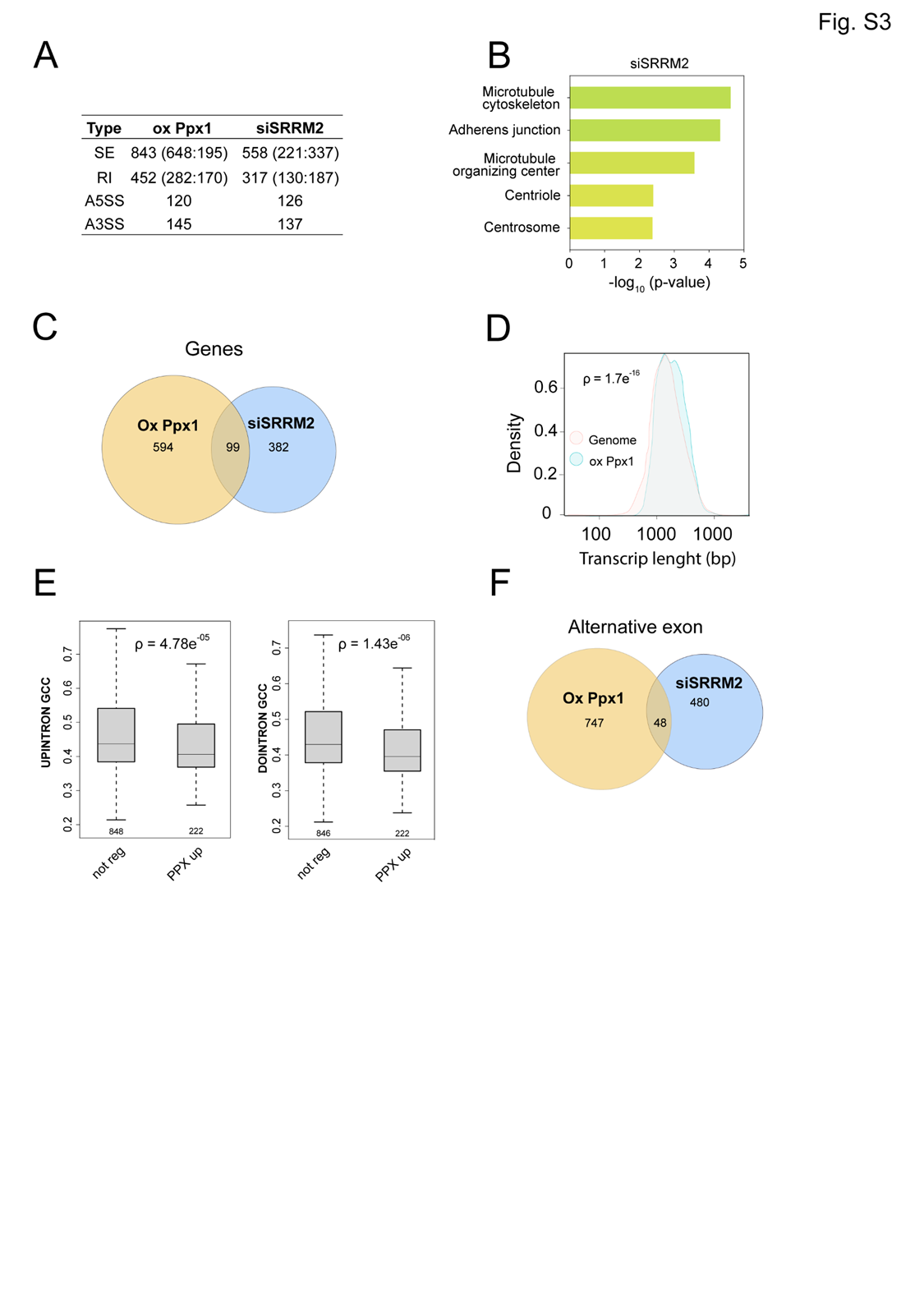

**Figure S3.** (A) Differentially spliced events in polyP and SRRM2 depleted cells. SE is for exon skipping, RI for intron retention, A5SS for alternative 5' splice-site and A3SS for alternative 3' splice-site. In brackets (inclusion: exclusion) events numbers. (B) GO subcellular function of exon skipping events in siSRRM2 cells. (C) Venn diagram showing overlapping genes in cells depleted for polyP and SRRM2. (D) Transcript length in polyP depleted cells vs standard genome. (E) GC content in the upstream intron of the splicing evens (upper panel). GC content in the downstream intron of the splicing event (lower panel). (F) Venn diagram of differentially spliced events in cells depleted for polyP and SRRM2.

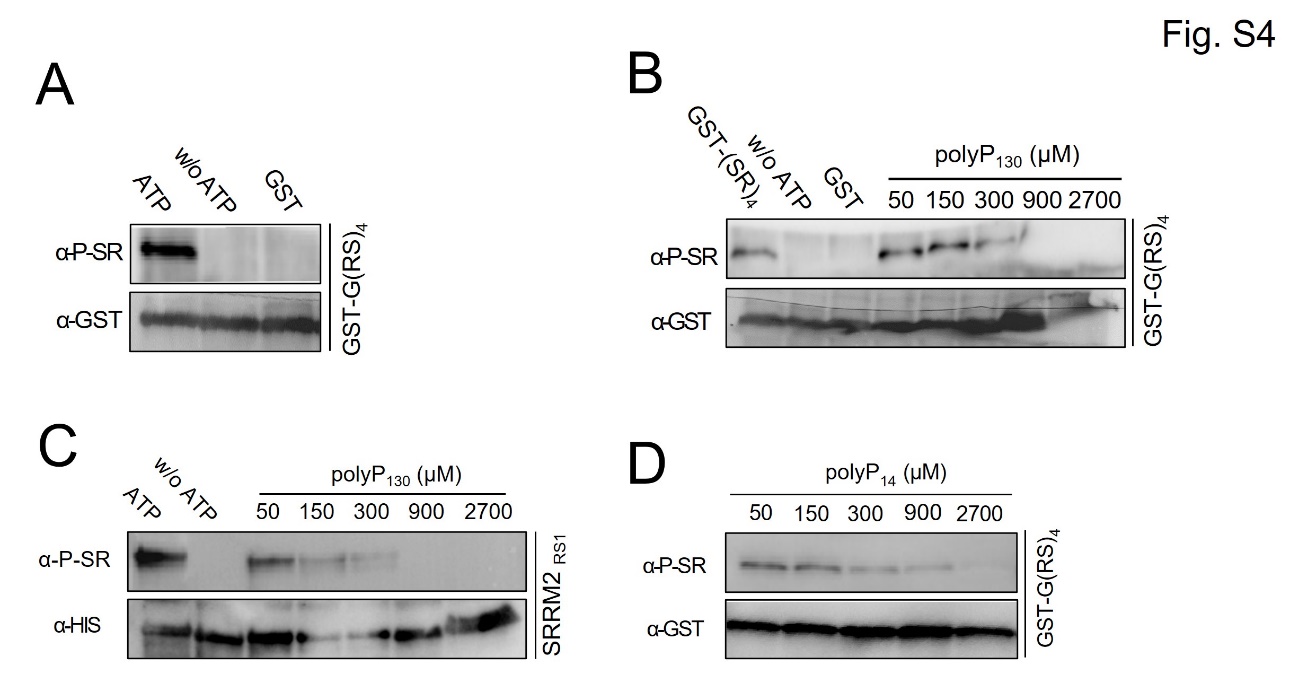

**Figure S4.** (A) CLK3 phosphorylates a peptide containing RS repeats. *In vitro* kinase assay of CLK3 obtained from insect cells incubated with a recombinant peptide containing 4 RS repeats. Phosphorylation was checked by western blot using a phospho-SR-specific antibody. GST antibody was used as load control and to discard mobility shifts due to protein-polyP interactions. (B, C) CLK3 kinase activity is inhibited by polyP. Same kinase assay as in A using as substrate the 4 repeats of the RS peptide (B) or a RS domain from SRRM2. As noted, growing concentrations of commercial 130-mer long polyP were included. (D) The same kinase assay as in A but using recombinant CLK3 produced in insect cells and including the noted amounts of 14-mer polyP.

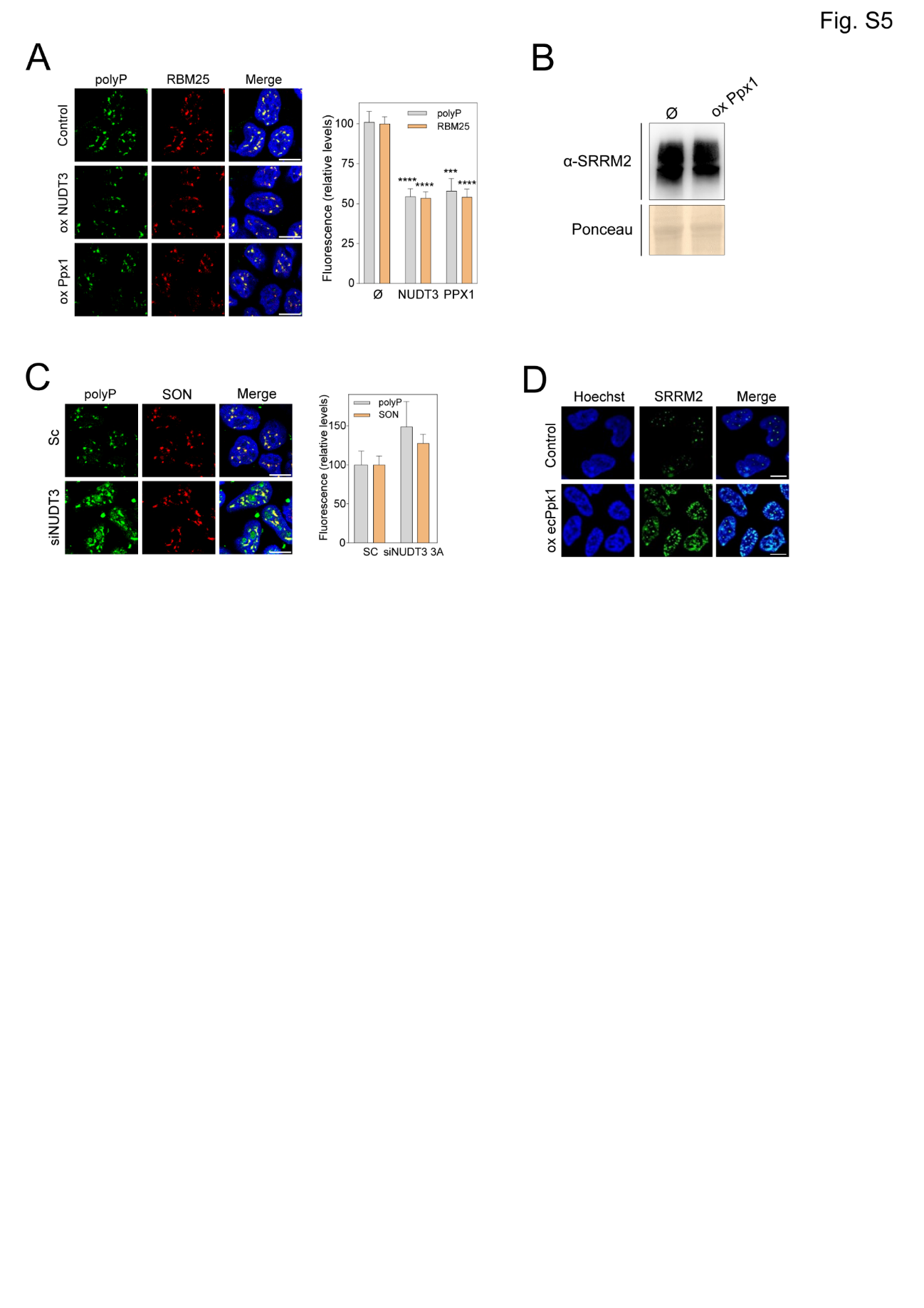

**Figure S5.** (A) RBM25 NS localization depends on polyP. Cells were transfected with empty pWPI, pWPI-NUDT3, or pWPI-Ppx1. After 48 h, polyP and RBM25 were quantified by immunofluorescence. A minimum of 500 cells per condition were analyzed. Mean intensity ± SEM of at least fifteen pictures of three independent experiments is shown. ***p < 0.001; ****p < 0.0001. (B) SRRM2 content in polyP-depleted cells. Cells were transfected with empty pWPI or pWPI-Ppx1. After 48 h, SRRM2 levels were assessed by western blot using specific antibodies. (C) Increasing polyP amount results in SON accumulation. Cells were transfected with scrambled or NUDT3-targeting siRNA. After 72 h, polyP and SON were quantified by immunostaining. A minimum of 500 cells per condition were analyzed. Mean intensity ± SEM of at least ten pictures of one experiment is presented. (D) Polyphosphate overloading enlarges NS. Ppk1 (*E coli* polyP synthase) overexpression in TREx-PPK1 cells was induced during 48 h by adding doxycycline. Representative image.
